# On the lipid dependence of bacterial mechanosensitive channel gating in situ

**DOI:** 10.1101/2024.01.22.576706

**Authors:** Madolyn Britt, Katsuhiro Sawasato, Elissa Moller, Gerald Kidd, Mikhail Bogdanov, Sergei Sukharev

## Abstract

For bacterial mechanosensitive channels acting as turgor-adjusting osmolyte release valves, membrane tension is the primary stimulus driving opening transitions. Because tension is transmitted through the surrounding lipid bilayer, it is possible that the presence or absence of different lipid species may influence the function of these channels. In this work, we characterize the lipid dependence of chromosome-encoded MscS and MscL in E. coli strains with genetically altered lipid composition. We use two previously generated strains that lack one or two major lipid species (PE, PG, or CL) and engineer a third strain that is highly enriched in CL due to the presence of hyperactive cardiolipin synthase ClsA. We characterize the functional behavior of these channels using patch-clamp and quantify the relative tension midpoints, closing rates, inactivation depth, and the rate of recovery back to the closed state. We also measure the osmotic survival of lipid-deficient strains, which characterizes the functional consequences of lipid-mediated channel function at the cell level. We find that the opening and closing behavior of MscS and MscL tolerate the absence of specific lipid species remarkably well. The lack of cardiolipin (CL), however, reduces the active MscS population relative to MscL and decreases the closing rate, slightly increasing the propensity of MscS toward inactivation and slowing the recovery process. The data points to the robustness of the osmolyte release system and the importance of cardiolipin for the adaptive behavior of MscS.

## Introduction

The mechanosensitive channel of small conductance, MscS, is one of the two major osmolyte release valves in most bacteria. It opens by membrane tension and curbs excessive osmotic gradients held by the cytoplasmic membrane, thereby adjusting turgor pressure within physiological limits. In most free-living bacteria, the low-threshold MscS channel works in tandem with the large-conductance and high-threshold mechanosensitive channel MscL, saving bacterial cells from lysis in the event of extreme media dilution such as in a sudden rain storm (1–3) or during an aquatic cycle of pathogens or commensals transmitted through fresh water (4).

Historically, MscS-like channels’ activities were the first observations of tension-activated channels in giant bacterial spheroplasts (5). These activities evoked by suction applied to the patch pipette were modulated by lipid-soluble amphipathic substances (CPZ, TNP), which indicated that these channels are sensitive to the distribution of forces in the surrounding lipids and suggested that these types of channels gate by tension in the lipid bilayer (6). Biochemical separation of solubilized E. coli membrane extracts and liposome reconstitution of individual fractions revealed that MscS and MscL are separate proteins that reliably produce native-like tension-activated currents when reconstituted in exogenous soybean lipids (7). This again strongly suggested that these channels are gated directly by tension transmitted through the lipid bilayer. Identification of MscL (8) and MscS (1) cloning opened broad opportunities to explore their mechanisms. The in vitro expression of MscL (8), as well as the reconstitution of affinity-isolated MscL (9) and MscS (10) in pure lipids, unequivocally confirmed that simply embedding these proteins into a generic lipid bilayer and stretching it with a 6-12 mN/m tension is sufficient to invoke channel activities with no additional components required (11–13). This mechanism of direct gating by force from lipids (FFL) has been conceptualized and reviewed (14–16).

The question of how different lipids may interact with the tension-activated channels and influence their function has been only partially clarified. MscL, with its characteristic ‘flattened’ open conformation (17), gated at lower tension in short-chain lipids (18,19), but gated at higher tensions in PE compared to PC liposomes, apparently due to sensitivity to the lipid headgroup composition (20). MscS, in contrast, was suggested to be less sensitive to the chemistry of the headgroups but was shown to be sensitive to the degree of unsaturation of aliphatic chains and the presence of cholesterol, which was related to the compressibility of the surrounding bilayer (21). The attempts to pinpoint specific lipids that may influence the gating were done in reconstituted proteoliposomes (22) and indicated that the presence of cardiolipin removes gating hysteresis in pressure-ramp experiments by increasing the closing rate. Cardiolipin has been independently implicated in the functions of energy-transducing and transport systems located in the inner membrane of bacteria and mitochondria (23–25). More recently, its requirement was shown for the stability of the protein secretion SecYEG/SecA system, which was strongly compromised in CL-lacking *E. coli* mutants (26).

Although, it was proposed that PG and CL are equally potent and interchangeable in many membrane-bound processes, *E. coli* cells lacking CL display multiple defects that would seriously compromise their viability outside of the laboratory. As in other studies in which CL has been reduced or depleted a compensatory rise in anionic PG was observed (25,27–29). Nevertheless, by taking advantage in an *E. coli* mutant fully devoid of CL we demonstrated recently that despite an absence of growth defects, co- and post-translational translocation of a-helical proteins across the inner membrane and the assembly of outer membrane β-barrel precursors were severely compromised in CL-lacking cells. *E. coli* CL-deficient strains display several deficiencies, which would limit their survival, vitality or adaptability under stressed conditions. The absence of CL activates the Rcs envelope stress response, which represses the assembly and production of flagella, disrupts surface adhesion and biofilm attachment, and reduces biofilm formation and growth (28,30)(Rowlett et al., 2017; Nepper et al., 2019).

Deficiencies in CL result in drastic and pleiotropic effects on E. coli’s cell envelope ultrastructure, maintenance of cell size and proper cell division (Rowlett et al., 2017).

In this work, we continue to characterize the lipid dependence of chromosome-encoded MscS and MscL in E. coli strains with genetically altered lipid composition. We use two previously generated strains that lack one or two major lipid species (PE, PG, or CL) and describe the engineering of a third strain that is highly enriched in CL and lacks PG. Using patch-clamp, we characterize the functional behavior of native mechanosensitive channels in these strains and quantify the relative tension midpoints, closing rates, inactivation depth, and recovery rate back to the closed state. In addition, we characterize the osmotic survival of lipid-deficient strains as the end result of the functional impact specific lipids on these channels. We find that the opening and closing behavior of MscS and MscL generally tolerate the absence of certain lipid species. The lack of cardiolipin (CL), however, visibly affects the propensity of MscS toward inactivation and slows down the process of recovery. The data points to the importance of cardiolipin for the adaptive behavior of MscS.

## Methods

### Bacterial strains

The ‘wild-type’ E. coli W3899 strain (31) was the parental strain for the below strains.

AL95 (pss93::kan lacY::Tn9) cannot synthesize PE. Therefore, it contains only acidic phospholipids PG and CL (45% and 50%, respectively) and phosphatidic acid, which makes up an inner membrane (IM) consisting of only negatively charged phospholipids (32,33). This strain was grown in an LB liquid medium supplemented with 50 mM MgCl2, which is required for viability.

UE54 (lpp2 Δara714 rcsF::mini-Tn10cam ΔpgsA::FRT-Kan-FRT) carries a null allele of the pgsA gene encoding the committed step to phosphatidylglycerol (PG) and cardiolipin (CL) biosynthesis making it devoid of PG and CL and containing at the stationary phase of growth about 95 mol% phosphatidylethanolamine (PE), 3 mol% phosphatidic acid (PA) and 2 mol% CDP-diacylglycerol (34) BKT12 (ΔclsA, ΔclsB, ΔclsC1::KanR) lacks all three cardiolipin synthase (cls) genes and does not synthesize CL (26,27,35).

The BKT12 (ΔclsABC) /pBAD30- ClsA G51A strain is the same as above but complemented with the plasmid carrying the G51A gain-of-function mutant of cardiolipin synthase ClsA. This construct is reported for the first time. To fully express plasmid-borne ClsA in BKT12 (ΔclsABC) /pBAD30- ClsA G51A was induced with 0.25% arabinose typically until OD_600_ reached 0.6.

### Construction of ClsA G51A mutant

ClsA is the major log-phase cardiolipin synthase in E. coli. The pBAD30-ClsA (EE) G51A mutant plasmid was generated from the pBAD30- ClsA (EE) template with QuikChange II Site-Directed Mutagenesis Kit (Agilent). Two residues, P48 and G51, were mutated using primers: clsA-P48A-5’ TTGATTATTTACATTCTGGCTTTAGTCGGAATTATTGCC clsA-P48A-3’ GGCAATAATTCCGACTAAAGCCAGAATGTAAATAATCAA clsA-G51A-5’ TACATTCTGCCGTTAGTCGCTATTATTGCCTATCTTGCC clsA-G51A-3’ GGCAAGATAGGCAATAATAGCGACTAACGGCAGAATGTA

The mutations were verified by automated DNA sequencing. The genes encoding ClsA with the P48A and G51A mutations were cloned in XbaI/HindII sites of commercial pBAD30 AmpR (p15A) under an arabinose-inducible promoter.

### TLC analysis of lipids

To determine the steady state phospholipid composition by radiolabeling, cells were uniformly labeled with 5 µCi/ml of [^32^P]PO_4_ after dilution to OD_600_ of 0.05 to initiate logarithmic growth or stationary growth after dilution of the overnight culture. Cells were harvested by centrifugation, and phospholipids were extracted by acidic Bligh Dyer procedure and analyzed after separation by one-dimensional TLC. ^32^P-labeled or non-radiolabeled phospholipids were resolved by thin layer chromatography (TLC) using HPTLC 60 silica gel plates (EMD, Gibbstown, NJ) activated and developed with a solvent consisting of chloroform:methanol:acetic acid (65:25:8 v/v) or (chloroform/ methanol/water/ammonium hydroxide [60:37.5:3:1

(vol/vol)]) as described previously (27,36). The spots were assigned to different E. coli phospholipids, aminophospholipids, and their derivatives based on TLC mobility (Rf values) of in-house standards. Radiolabeled lipids were visualized and quantified using a Typhoon FLA 9500 Imager (GE). Stored images were processed and quantified using ImageQuantTM software. Phospholipid content is expressed as mol% of total phospholipid (correcting for two phosphates per molecule of CL) based on the intensity of the captured signal on the Phosphor screen generated by the radiolabeled spots on the TLC plate. The results presented are representative of three or more determinations.

### Electrophysiology

Giant spheroplasts for patch-clamp experiments were generated as previously by inducing filamentous growth with cephalexin followed by lysozyme digestion in the presence of EDTA (5). The exception was the PE-containing UE54 strain that did not grow in cephalexin. Its filaments were grown in carbenicillin (0.3 mg/ml) instead. Recordings were made on excised inside-out patches in a symmetric buffer containing 400 mM sucrose, 200 mM KCl, 50 mM MgCl_2_, 5 mM CaCl_2_, and 5 mM HEPES at pH 7.4. Standard borosilicate glass pipettes (Drummond) were pulled to a bubble number of 4.8-5 (ref) to control the size. Currents were recorded using an Axopatch 200B amplifier (Molecular Devices). Negative pressure stimuli applied to the pipette were programmed using the Clampex software (Molecular Devices) and applied through a high-speed pressure clamp apparatus (HSPC-1; ALA Scientific) with modified suction pumps. Data acquisition was performed with the PClamp 10 suite (Molecular Devices).

Characterization of channel populations in each patch typically began with recordings of current responses under symmetric 1-s linear pressure ramps at +30 mV (pipette voltage). The shapes and amplitudes of two-wave responses reflecting separate MscS and MscL populations provided midpoint and saturating pressures for each population.

Additional protocols were used to extract kinetic and thermodynamic parameters associated with opening, closing, and inactivating transitions. Closing rate pulse-step protocols were performed to determine tension-dependent closing rate parameters. Inactivation “comb” protocols in which channels closing under tension are periodically probed with saturating pulses to determine what portion of the population remains tension-sensitive (37) were employed to monitor the rate of inactivation. Recovery from inactivation pulse-step-pulse protocols were also done by pulsing at saturating tensions to reveal the full population, holding at the midpoint pressure to encourage channels to close and inactivate, and then pulsing again with a series of saturating pressure pulses to extract the rates of recovery from the inactivated to the resting state. Experiments were performed at ±30 and ±60 mV to test the voltage-dependent character of state transitions.

### Osmotic survival

Osmotic viability was determined by counting colony forming units (CFUs) /mL after osmotic shock from 1200 mOsm down to either 600 or 200 mOsm media. Cells were sub-cultured 1:100 from standard overnight cultures in Luria Bertani (LB) media into XhiLB (LB supplemented with NaCl to final osmolarity 1200 mOsm) and grown to early logarithmic phase (OD_600_ of ∼0.25). The BKT12 strain carrying the clsA G51A :: pBAD plasmid was induced with 0.25% arabinose for 1.5 hr before osmotic dilution. Down-shock was imposed by rapidly pipetting 50 uL of growth culture into 5 mL of 1200 mOsm LB (control), 600 mOsm LB, or 200 mOsm LB. Samples were incubated at room temperature for 15 minutes and then diluted 1:50 into the corresponding shock media. Cells were then plated in duplicate, and colonies were manually counted the following day. The fraction of osmotic survival was determined by normalizing cell counts relative to the un-shocked 1200 mOsm control. Experiments were performed independently at least 3 times, and the error shown is the standard deviation between these trials.

### Statistical analysis

All values are reported as N measurements’ mean and standard deviation (mean ± SD).

## Results

### The lipid distribution in the strains used for mechanosensitive channel characterization

The TLC patterns of separated lipids obtained from the four previously generated strains are shown in Fig. 1. W3899 is the wild-type strain with respect to all components of glycerophospholipid biosynthetic pathways and lipid composition. Phosphatidylethanolamine (PE), phosphatidylglycerol (PG), and cardiolipin (CL) comprise about 75%, 20%, and 5 % of the total phospholipids, respectively (Fig. 1A, lane 1).

**Fig. 1.**
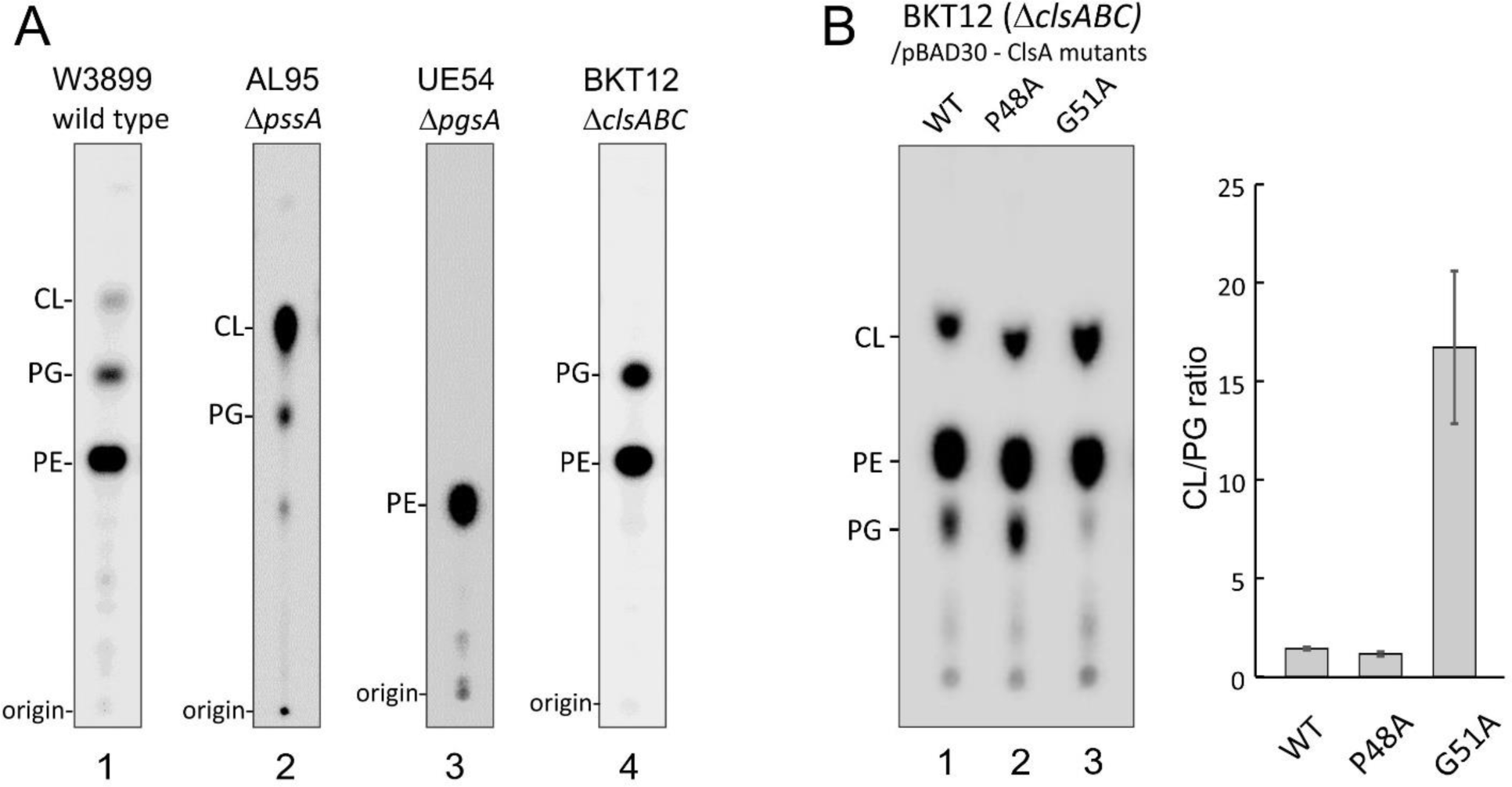
Lipid profiles of E. coli mutants with altered lipid compositions. The strains are denoted above the lanes. Uniformly 32P labeled phospholipids were extracted and analyzed after separation by one-dimensional TLC on boric acid-impregnated silica gel plates in solvent consisting of chloroform:methanol:acetic acid (65:25:8 v/v) (A) or in chloroform/ methanol/water/ammonium hydroxide [60:37.5:3:1 (vol/vol)] (B). (A) The lipid patterns in the four existing strains. (B) The lipid profiles in the BKT12 strain expressing WT ClsA (lane 1), the P48A (lane 2), and G51A (lane 3) mutants from pBAD30 plasmid. All versions were equally induced by 0.25% arabinose for 2 hrs. The panel on the right shows average CL/PG ratios obtained by TLC densitometry from three independent experiments, and the error bars represent S.D.

The AL95 strain (ΔpssA) cannot synthesize PE and, therefore, contains only acidic phospholipids PG and CL (45% and 50%, respectively) and about 5% of phosphatidic acid (Fig. 1A, lane 2).

UE54 carries a null allele of the pgsA gene encoding the committed step to PG and CL biosynthesis, making it devoid of PG and CL and containing at the stationary phase of growth about 95 mol% phosphatidylethanolamines (PE), 3 mol% phosphatidic acid (PA) and 2 mol% CDP-diacylglycerol (Fig. 1A, lane 3).

To study the role of CL in the functioning of mechanosensitive channels, we utilized the E. coli CL-deficient mutant BKT12 that lacked all three CL synthases (ΔclsABC1). The BKT12 mutant is entirely devoid of CL and utilizes PG as the primary anionic lipid species (Fig. 1A, lane 4).

### Generation of the cardiolipin-enriched strain

ClsA is the main log-phase cardiolipin synthase in E. coli that reversibly links two PG molecules producing one CL (27), thereby maintaining the ratio of the two anionic species during the active growth phase. As shown in Fig. 1B, the re-expression of WT ClsA from the pBAD plasmid restores the presence of CL in the BKT12 strain, but the molar ratio of different lipid species in this complemented strain is somewhat different from W3899 (WT). Preparation of gain-of-function or partial loss-of-function ClsA mutants with attenuated activity would become a convenient tool to regulate the ratio of anionic lipids and study functional consequences.

The structure of the 486-amino acid ClsA enzyme is still unavailable, but multiple models, including the one generated by alphafold2 (https://www.rcsb.org/structure/AF_AFP71040F1), predict a single N-terminal transmembrane anchoring domain followed by a cytoplasmic 430-aa catalytic domain. The transmembrane domain begins with Y5 on the periplasmic side and apparently ends with a positive cluster K30-R31-R32 on the cytoplasmic side forming a turn. After that cluster, the chain is predicted to plunge back into the membrane, forming the second short helix, which breaks at P48. The chain then emerges from the membrane to continue into the catalytic domain through a flexible hinge at G51. We argued that the dynamics and proximity of the catalytic domain to the bilayer surface may depend on the helical break and hinge positions in the accordance with established the relationship between Gly and Pro close spacing and conformational properties of a- helix (38).

The alanine substitutions at positions P48 and G51 had different effects, as shown by the relative density of PG and CL spots in the respective strains (Fig. 1B, lanes 2 and 3). The P48A substitution had no effect when compared to WT, whereas the G51A mutant plasmid showed a decisive gain-of-function phenotype, increasing the CL-to-PG ratio about 15-fold. We should note that the condensation of PG by ClsA is energy independent, and the G51A mutation in the second membrane-spanning domain makes ClsA hyperactive. This product of this construct, which almost entirely converts PG into CL, was then used to complement the CL-less BKT12 strain and produce the CL-rich BKT12 clsA G51A :: pBAD strain to study mechanosensitive channels and osmotic survival.

### Patch-clamp characterization of MS channels in individual lipid-deficient strains using pressure ramps

The parental strain W3899 (WT) readily formed filaments in the presence of cephalexin and produced patchable spheroplasts. A patch formed by our standard 1.3-1.4 um pipette (4.8-5 bubble number) subjected to a linear pressure ramp produces a characteristic two-wave current trace (Fig. 2A). The amplitude of the first wave, corresponding to the activation of the low-threshold channel population, looks uniform and likely contains primarily MscS. The second wave represents the activity of MscL on the background of a fully saturated MscS population. The numbers of channels per patch and the absolute midpoint pressures are presented in Fig. 3, and their ratios are presented in Table 1.

**Fig. 2.**
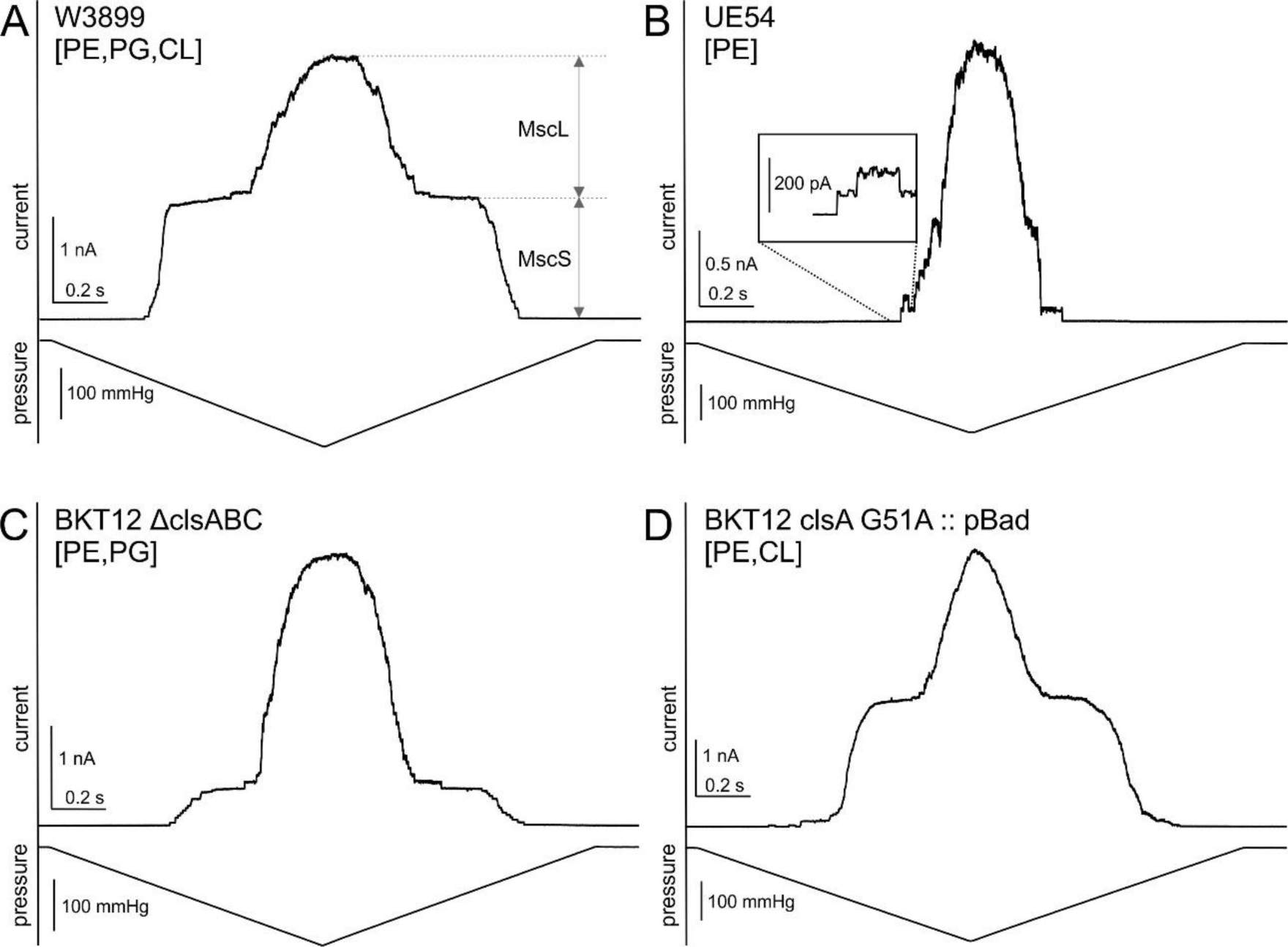
Representative current responses to pressure ramps for the four strains. The two waves of MscS and MscL channel activities are denoted in panel A for W3899 (WT). The UE54 strain (B) in 50% of patches exhibits no MscS current; in some patches, in addition to MscS and MscL, it shows a small number of early-activating low-threshold channels. Traces for the BKT12 ΔclsABC strain (C) show a lower number of MscS compared to the same strain complemented with the clsA G51A plasmid (D). The statistics of the channel content (Fig. 3B, Table 1) show differences between the strains.

**Fig. 3.**
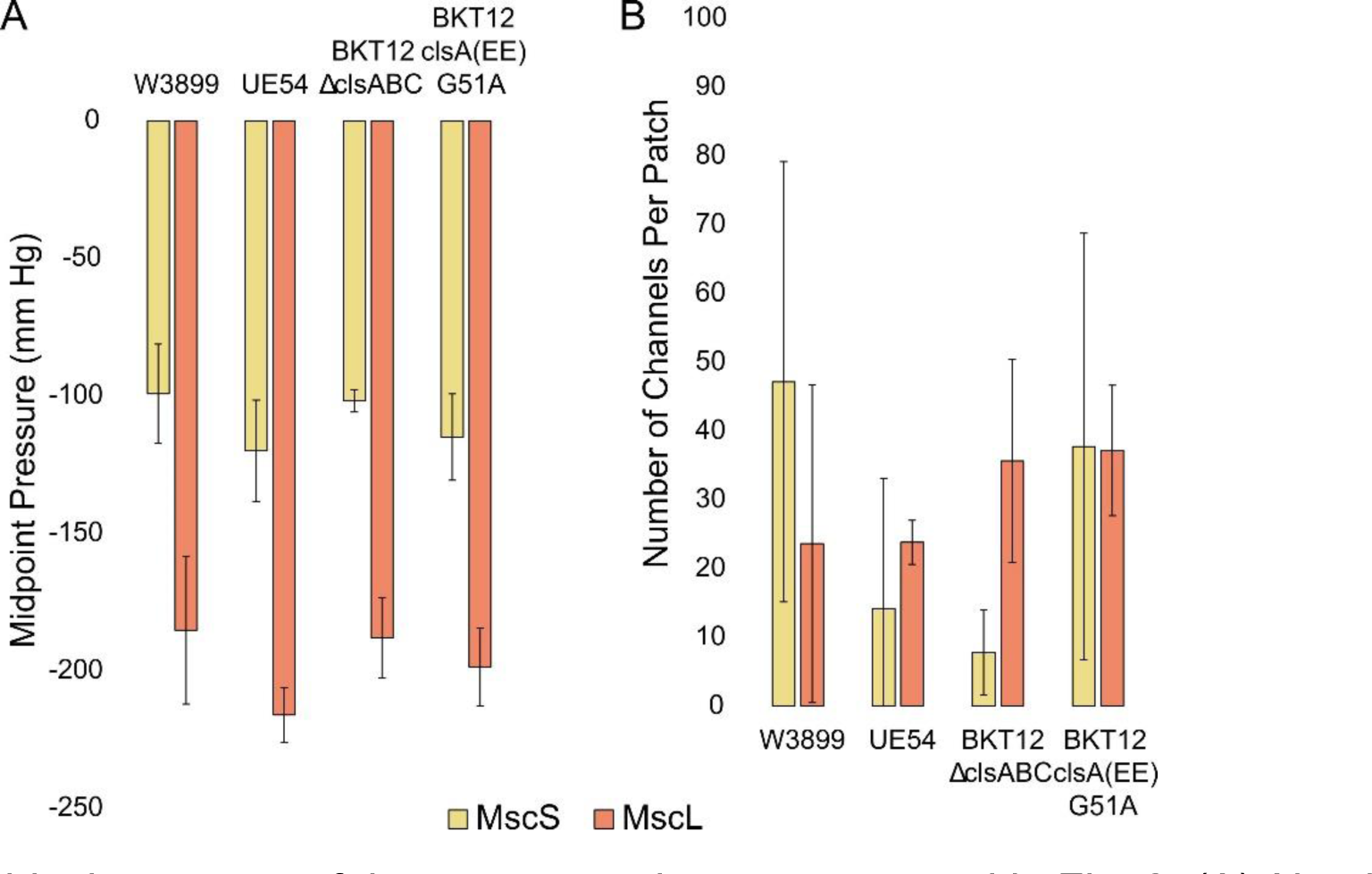
A graphical summary of the ramp experiments presented in Fig. 2. (A) Absolute values of midpoint pressures for the four strains indicate that there was no systematic simultaneous deviation of stimuli that would produce a similar P0.5MscS/ P0.5MscL ratio. (B) The numbers of MscS and MscL channels per patch suggest that the MscS activities tend to be lower without CL.

**Fig. S1.**
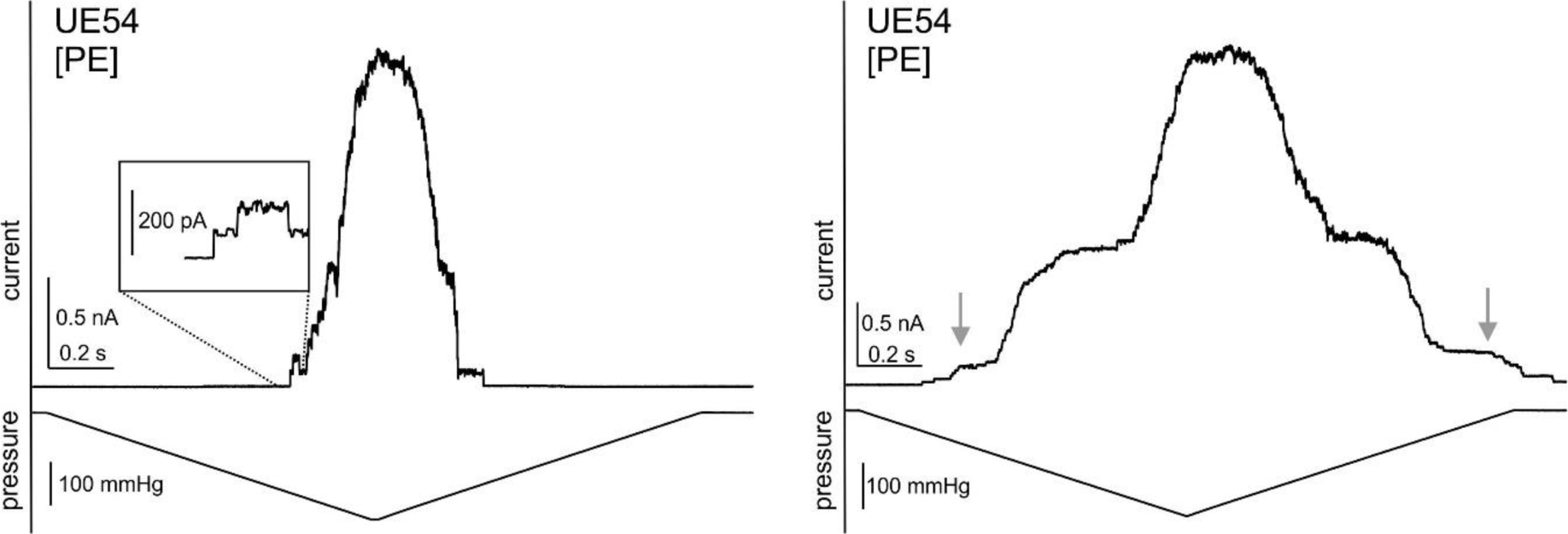
Illustration of ramp currents recorded on two consecutive patches. The first patch (left) has no MscS activities, whereas the second patch (right) shows the MscS population and, in addition, a smaller population of low-threshold mini-channels seesn as a small shoulder at the foot of the trace, on both ascending and descending legs of the pressure ramp.

**Table 1.** The activating pressure midpoints for MscS and MscL populations, their ratios, and the numbers of channels per patch based on ramp responses for the four strains.

|  | **P<sub>0.5</sub>MscS** | **P<sub>0.5</sub>MscL** | **P<sub>0.5</sub>MscS:**<br>**P<sub>0.5</sub>MscL** | **MscS**<br>**Midpoint**<br>**Tension**<br>**(mN /m)** | **# MscS**<br>**channels** | **# MscL**<br>**channels** |
| --- | --- | --- | --- | --- | --- | --- |
| **W3899**<br>**[PE,**<br>**PG,CL]** | -99 ± 18 | -185 ± 27 | 0.55 ± 0.07 | 7.7 ± 0.9 | 47 ± 32 | 24 ± 23 |
| **UE54**<br>**[PE]** | -107 ± ?? | -216 ± 10 | 0.51 ± ?? | 6.6 ± ?? | 18 ± 25 | 24 ± 3 |
| **BKT12**<br>**ΔclsABC**<br>**[PE,PG]** | -102 ± 4 | -186 ± 17 | 0.59 ± 0.04 | 7.9 ± 0.5 | 19 ± 15 | 32 ± 16 |
| **BKT12**<br>**pBad**<br>**[PE,CL]** | -115 ± 16 | -199 ± 14 | 0.58 ± 0.06 | 7.9 ± 0.8 | 38 ± 31 | 37 ± 9 |

The UE54 (ΔpgsA) strain containing primarily (only) PE did not grow in cephalexin at all and only poorly in carbenicillin. As a result, the spheroplasts were small, and the preparations were barely patchable. Yet, obtaining several patches and recording complete ramp responses was possible. Surprisingly, in the almost complete absence of negatively charged lipid species (no PG or CL), the ratio of pressure midpoints were similar to WT. While the absolute number of MscL channels per patch was the same as in WT, the number of MscS channels was lower. (Fig. 3B and Table 1A). In some UE54 patches, in addition to MscS and MscL, the UE54 strain shows a small number of early-activating low-threshold channels visible at the foot of the trace (Fig. S1, supplement) on both ascending and descending legs of the pressure ramp.

The BKT12 ΔclsABC and UE54 strains with altered content of negatively charged lipids (PG and CL) both show a lower proportion of MscS current and a higher fraction of MscL compared to WT (Fig. 3B). The BKT12 ΔclsABC strain lacks all three cardiolipin synthases and has no CL, only PG. The BKT12 clsA G51A :: pBAD strain, carrying a gain-of-function *clsA* version on a plasmid that converts all available PG into CL, has PE and CL. Induction of this plasmid somehow equalizes the numbers of MscS and MscL per patch (Fig. 3B). With these moderate deviations, none of the three mutant strains show any drastic deviation from WT. Regarding the activation midpoint, the strains also show consistency. Only the PE-containing UE54 seemed to show a slightly lower midpoint for MscS. We should note that in CL-free UE54 and BKT12 ΔclsABC strains, about 50% of patches had no MscS channels, only MscL.

### Determination of the spatial parameters and rates for MscS closing

In the following sections, we focus on the properties of MscS only. Even though each patch contained two populations, MscS and MscL, it was possible to record and characterize only MscS because the threshold of MscL activation is sufficiently far above the saturating pressure for MscS. Occasionally, we saw a small admixture of MscL activities, which were recognized, and the pressure protocols were adjusted accordingly to avoid extra activity.

The closing rates of MscS channels were determined with the pulse-step protocol. The short saturating pressure pulse opens the entire population in the beginning, then the pressure is dropped to a series of intermediate levels and the current relaxation is recorded. Fitting the relaxation curves with monoexponential functions produces plots of the closing time constants or kinetic rates as a function of step pressure. Extrapolation to zero pressure estimates the intrinsic closing rate. Among the three preparations that cooperated with this protocol (W3899, BKT12 and BKT12 clsA G51A :: pBAD), the slowest closing rate was for the CL-free BKT12 strain (Table 2).

**Table 2.**
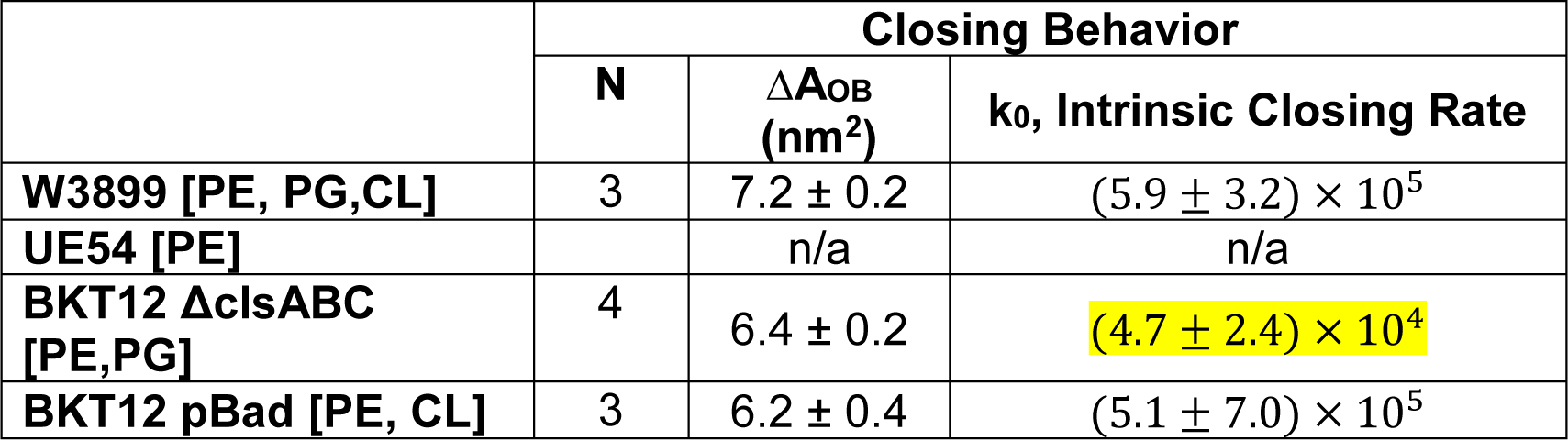
The spatial and kinetic parameters of the MscS closing transitions for the four strains. ΔA_OB_ is the distance from the bottom of the open-state well to the transition barrier (Fig. 4D), and k_0_ is the intrinsic closing rate determined by extrapolating experimental closing rates to zero tension (Fig 4C).

**Table 3.** The characteristic times of MscS inactivation at different voltages in the four strains.

|  | Inactivation Tau (s) |  |  |  |
| --- | --- | --- | --- | --- |
|  | -60 mV | -30 mV | +30 mV | +60 mV |
| **W3899 [PE, PG, CL]** | $2.2 \pm 1.5$ (4) | $7.8 \pm 6.3$ (5) | $6.3 \pm 1.2$ (4) | $6.0 \pm 0.6$ (3) |
| **UE54 [PE]** |  |  |  |  |
| **BKT12 ΔclsABC [PE,PG]** | $2.2 \pm 0.8$ (2) | $5.6 \pm 3.0$ (4) | $7.0 \pm 4.0$ (3) | $6.7 \pm 1.9$ (2) |
| **BKT12 pBad [PE,CL]** | $1.4 \pm 0.5$ (2) | $6.7 \pm 5.9$ (4) | $10.5 \pm 8.9$ (4) | $6.4 \pm 6.9$ (2) |

### The MscS inactivation rates

In the next experiment, we studied the inactivation rate with the previously described ‘comb’ pressure protocol (ref). This protocol includes a prolonged (10 or 20 s) conditioning pressure step with the amplitude near the activation midpoint interrupted with several saturating test pulses. The first test pulse activates the total population, indicating the available active channels. The moderate tension clamped by the conditioning step slowly drives the population to the inactivated state, which is tension insensitive, and the following test pulses indicate the fraction of adapted channels that are still available to open. In Fig. 4A, the tips of current responses to test pulses (arrows) indicate the kinetics of inactivation of MscS in the WT strain. The recordings at two different voltages show that the process of inactivation is sped up at depolarizing 30 mV (-30 mV in the pipette) compared to hyperpolarizing 30 mV. The current traces recorded from UE54 (Fig. 4B) look different, showing that the relaxation of the current from the peak value at the test pulse down to the intermediate level is much slower.

**Fig. 4.**
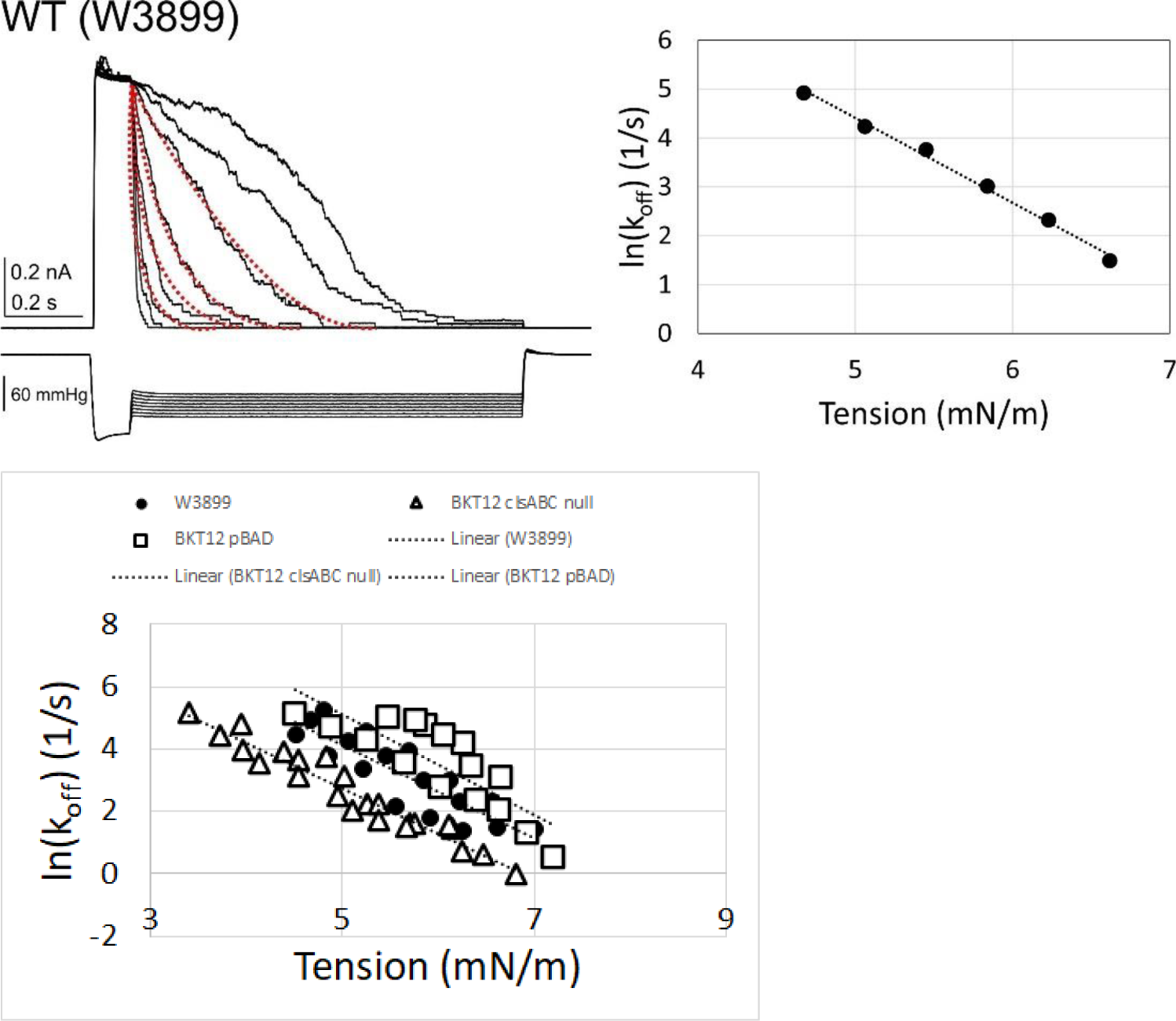
The determination of the closing rate using the pulse-step protocol. We need to add traces just for WT (A) and BKT12 ΔclsABC (B) for comparison, not for all four. We need the *ln*K_off_(gamma) plot for all of them (C). A schematic representation of the energy profile along the opening pathway where the reaction coordinate (x-axis) is the in-plane area of the MscS protein complex in the membrane (D).

**Fig. 4.**
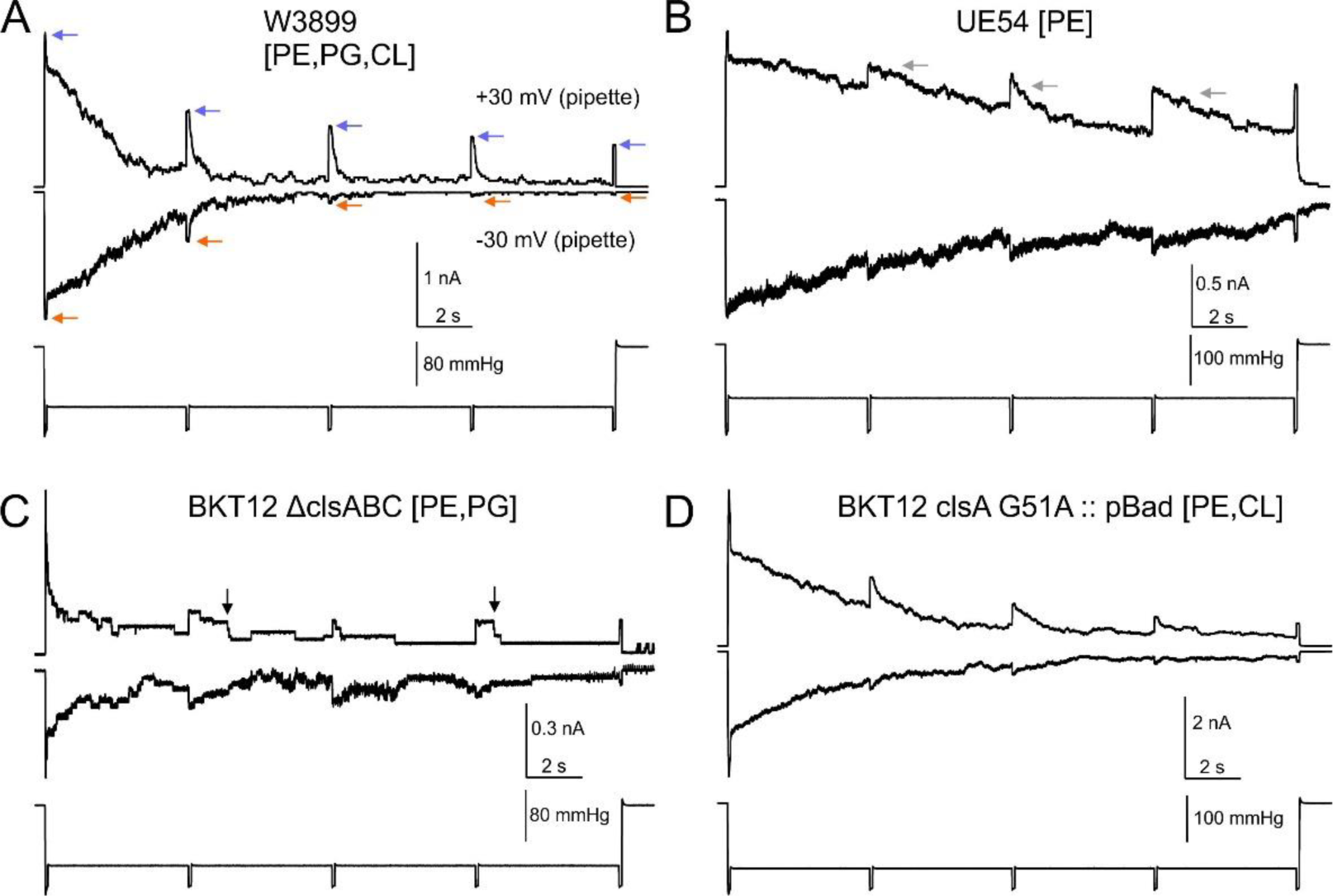
The inactivation rate of the four strains studied with the ‘comb’ protocol. The tips of current responses in WT (A) reflect the remaining active fraction of channels at hyperpolarizing (blue arrows) and depolarizing (orange arrows) 30 mV. (B) The responses in UE54 show slow current relaxation (grey arrows). The trace recorded from the BKT12 strain (C) illustrates a smaller channel population and a delayed closure. The plasmid-complemented BKT12 strain (D) shows a smoother closing.

The BKT12 trace (panel C) also shows open channels ‘lingering’ for much longer after each test pulse, indicating slow closure in the absence of CL. Consistent with the data on the closing rates (Table 2), complementing this strain with the hyperactive clsA enzyme tends to restore the closing kinetics. This suggests that negatively charged species, especially CL, help maintain a normal closing rate.

### The kinetics of MscS recovery from inactivation

The protocol for visualizing the recovery process consisted of the initial test pulse revealing the full population of active channels with a subsequent conditioning step of intermediate tension, during which the population gradually inactivates, followed by a train of test pulses that visualize recovery (Fig. 5).

**Fig. 5.**
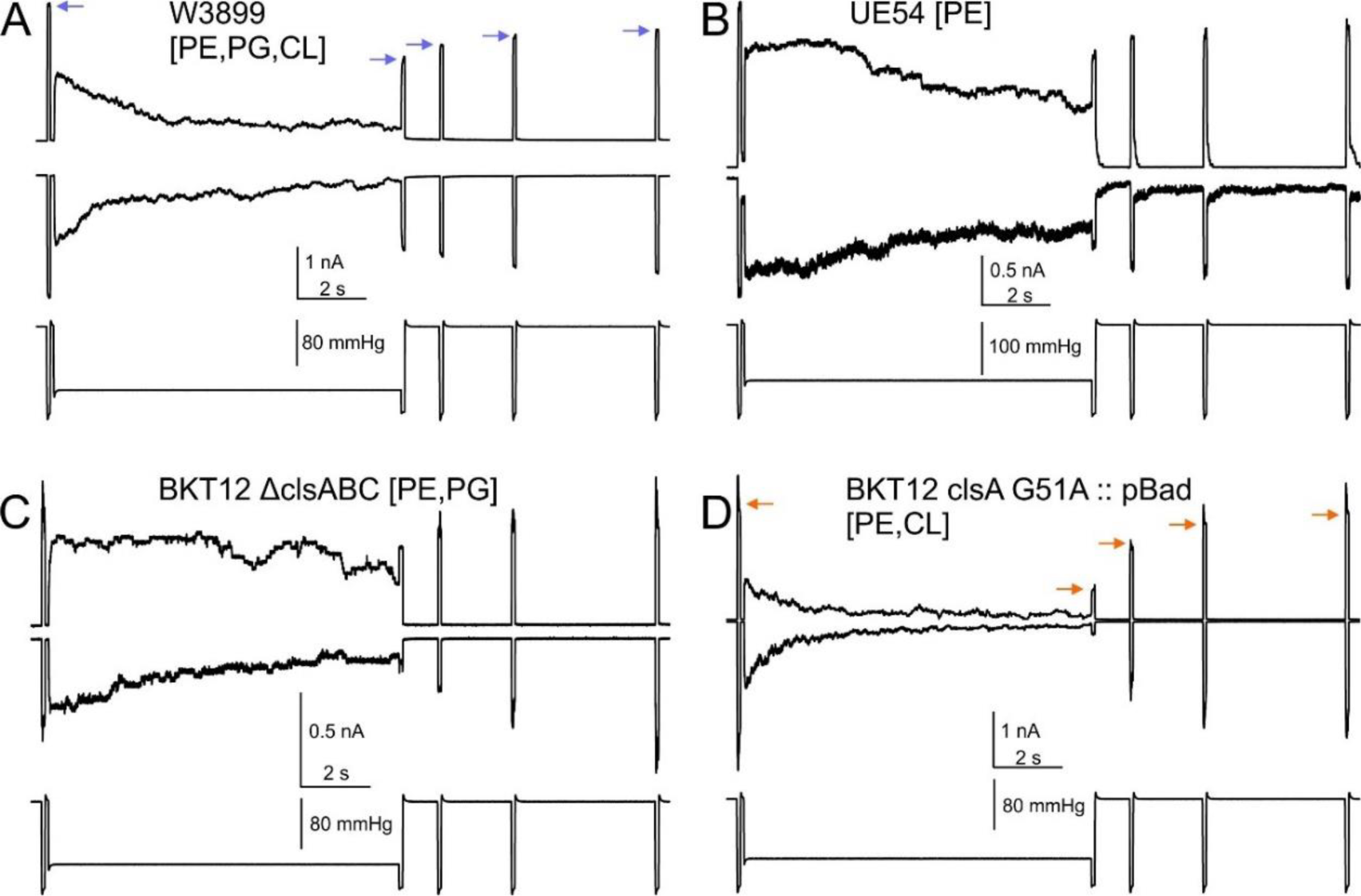
Characterization of inactivation and recovery of MscS in the four strains. The current amplitude at the tips of test pulse responses (arrows in panel A) is plotted against time and fitted to the monoexponential function to extract the characteristic recovery time.

**Fig. 5.**
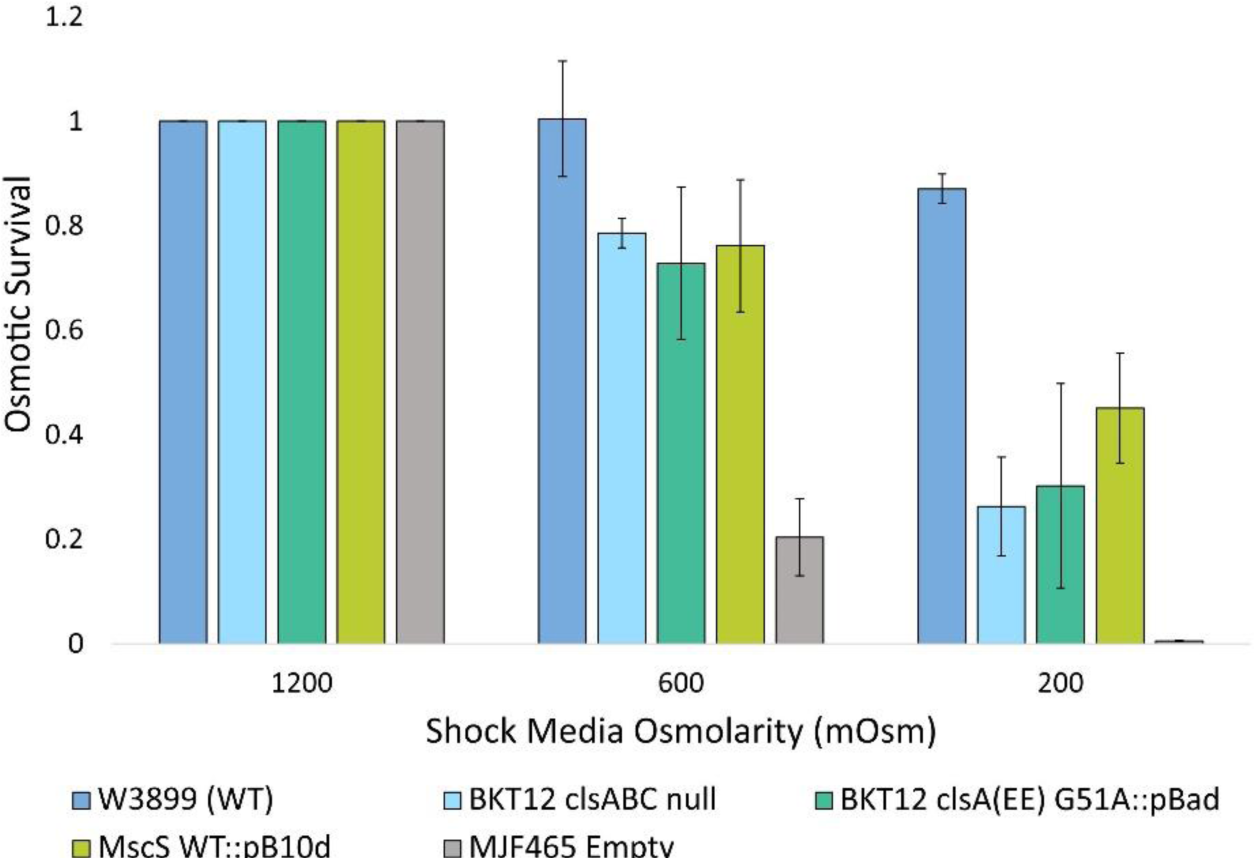
Osmotic survival of lipid-depleted strains under 1200 - 600 and 1200 - 200 mOsm shocks scored against the parental W3899 strain and previously characterized MJF465 strain protected by expression of MscS. The empty MJF465 (*ΔmscS ΔmscK ΔmscL*) strain served as the negative control.

The ratio of the current amplitude in response to the second test pulse at the end of the conditioning step to the initial test pulse response gives us the extent of inactivation during that step. The fitting of normalized current amplitudes indicated by arrows in Fig. 5A with a monoexponential function gave us values of tau (characteristic time) of recovery. The extent of inactivation and the recovery times determined at four different voltages are presented in Table 4. The recovery from inactivation was slightly slower in the CL-depleted BKT12 ΔclsABC strain.

**Table 4.** The dependences of the extent of inactivation on voltage and the characteristic recovery time.

|  | Extent of Inactivation (%) |  |  |  |
| --- | --- | --- | --- | --- |
|  | -60 mV | -30 mV | +30 mV | +60 mV |
| **MscS WT [PE, PG, CL]** | 97 ± 5 (6) | 74 ± 21 (10) | 41 ± 26 (10) | 29 ± 25 (8) |
| **W3899 [PE, PG, CL]** | 97 ± 2 (4) | 65 ± 26 (5) | 35 ± 8 (4) | 46 ± 26 (5) |
| **UE54 [PE]** |  | 42 ± ?? (1) | 29 ± 12 (2) |  |
| **BKT12 ΔclsABC [PE,PG]** | 92 ± 8 (4) | 80 ± 10 (7) | 51 ± 13 (8) | 51 ± 12 (7) |
| **BKT12 pBad [PE,CL]** | 97 ± 1 (3) | 78 ± 175 (8) | 56 ± 21 (9) | 48 ± 19 (3) |

|  | Recovery from Inactivation Tau (s) |  |  |  |
| --- | --- | --- | --- | --- |
|  | -60 mV | -30 mV | +30 mV | +60 mV |
| **MscS WT [PE, PG,CL]** | 2.1 ± 0.7 (6) | 1.3 ± 0.5 (10) | 1.1 ± 0.4 (7) | 1.3 ± 0.6 (8) |
| **W3899 [PE, PG,CL]** | 1.5 ± 0.45 (3) | 2.0 ± 0.5 (5) | 1.7 ± 0.2 (4) | 1.0 ± 0.2 (4) |
| **UE54 [PE]** |  | 1.2 ± ?? (1) | 0.8 ± 0.5 (2) |  |
| **BKT12 ΔclsABC [PE,PG]** | 3.3 ± 1.5 (4) | 2.4 ± 0.9 (6) | 3.7 ± 2.4 (9) | 1.0 ± 0.3 (6) |
| **BKT12 pBad [PE,CL]** | 5.5 ± 6.2 (2) | 2.8 ± 1.3 (7) | 2.0 ± 0.8 (8) | 1.4 ± 0.5 (3) |

### Osmotic survival of strains with altered lipid composition

The osmotic survival experiments were performed to assess the functionality of these channels in actively metabolizing cells. First, cells were grown in an XhiLB medium supplemented with NaCl to 1200 mOsm. Upon reaching OD 0.3, the early log cells were diluted into either 600 or 200 mOsm media and plated. The data presented in Fig. 5 shows that all mutant strains are compromised regarding osmotic survival. As positive controls, we used the parental W3899 (WT) strain as well as the previously characterized MJF465 triple mutant (ΔmscS ΔmscK ΔmscL) strain (1) empty and without any complemented channels and (1) expressing MscS from the pB10d plasmid (ref). Interestingly, the survival of lipid mutants was on par with that of the well-protected MscS-expressing strain (ref). The survival of the parental strain was the highest.

The UE54 (ΔPgsA) strain containing primarily PE did not grow in XhiLB (1200 mOsm), so we do not present its survival data. (AL95 strain to be added to the survival plot).

## Discussion

This work presents the first phenomenological characterization of mechanosensitive channel activities in three *E. coli* mutant strains with drastically altered lipid compositions. The differences in lipid profiles presented in Fig.1 are not subtle. In many strains, the content of specific lipids types is essentially ‘all’ or ‘none.’ With the exception of AL95 (no PE), in which the majority of lipids are charged, and the strain requires 50 mM MgCl_2_ for survival and propagation, all other strains did not show substantial growth defects in the LB medium, i.e., under standard laboratory conditions. Increasing the osmolarity to 1200 mOsm suppressed the growth of UE54, which contains only PE. This strain, devoid of PG and CL, also did not form filaments in the presence of cephalexin, which can be explained by altered peptidoglycan-synthesizing machinery since cardiolipin is required for many secretion and transport mechanisms (25,26,39). The oppositely skewed balance between PG and CL in BKT12 and BKT12 clsA G51A :: pBad strains did not compromise the growth, but the complete absence of CL in BKT12 made this strain more vulnerable to osmotic down shock compared to the other two mutant strains tested.

Looking at the patch-clamp data collected in lipid mutants, we can say that both MscS and MscL are generally robust and relatively lipid-insensitive channels. While the lipid composition changed drastically, neither the pipette pressures at which we observed the activation midpoints (Table 1A) nor the ratios of P0.5_MscS_/P0.5_MscL_ exhibited any systematic changes. We only observed a depression in the number of active MscS channels in two strains (UE54 and BKT12) devoid of CL (Table 1B). In about 50 % of BKT12 patches, we observed no MscS channels (only MscL, Fig. 2B). The problem of patch stability in UE54 precluded extensive characterization of the closing kinetics; however, in the other CL-free BKT12 strain, the intrinsic closing rate was one order of magnitude slower than in the complemented BKT12 clsA G51A :: pBad strain or WT. This points to the visible effect of cardiolipin on MscS kinetics: CL makes the channel close faster. This is highly consistent with the previous observation published by Ridone (22) that 10% of cardiolipin added to MscS-containing azolectin proteoliposomes increases the activation midpoint and removes gating hysteresis in symmetric ramp responses. This effect was interpreted as destabilizing the open state and stabilizing the closed state of MscS by CL in the azolectin environment. In the presence of CL, both MscS and MscL exhibited increased activation thresholds. However, we did not observe this effect systematically. CL also appears to affect the inactivation and recovery processes. Inactivation tends to slow down in CL-rich BKT12 clsA G51A :: pBad strain, whereas the reverse recovery process slows down in the CL-devoid BKT12 strain.

Cardiolipin seems to affect MscS by stabilizing its active state. What is unique about CL? It is a doubly-charged anionic lipid that tends to bind to sites in proteins organized around closely spaced pairs of arginines or lysines, preferably in the cytoplasmic leaflet of the cytoplasmic membrane (40). MscS, possessing 7 pairs of closely spaced arginines (R46 and R74) facing the phosphates of the inner leaflet (30), appears to have a probable location where CL might be enriched. As a dimer of two PGs, CL has four acyl chains, which makes it unique in terms of its bulkiness and its inability to permeate interhelical pockets and crevices in proteins. Its small headgroup and bulky aliphatic side also create an imbalanced lateral pressure profile with a maximum in the aliphatic core, exerting significant pressure on neighbors in the middle of the membrane. With these two respects (size and pressure distribution), cardiolipin must be a robust stabilizing factor providing inner ‘filler’ for proteins with irregular lipid-facing surfaces.

MscS’s interface with lipids is notoriously irregular. The stable splayed (conical) conformation observed in many structures (41,42) shows deep crevices often populated with regular two-acyl-chain lipids (PE or PG). These lipids separate the peripheral TM1- TM2 helices from the gate-forming TM3 barrel. For this reason, the splayed conformations with intercalated lipids likely represent the inactivated state. We presume that cardiolipin will not fit in the crevices, and this way, it will serve as a stabilizing factor for the more compact closed ready-to-open state. In contrast to many systems where cardiolipin is believed to be a structural part (39,43), we presume that not specific binding, but rather a presence around the protein and exclusion from the pockets provides the stabilizing effect countering massive inactivation.

## Acknowledgements

This work was funded in part by National Institutes of Health grants R01-GM121493 (to M.B.), R01AI135015 (to S.S.), and by the National Science Foundation Graduate Research Fellowship Program under grant No. DGE 1840340 to E.M.

## Notes

### Competing Interest Statement

The authors have declared no competing interest.

